# Source imaging of high-density visual evoked potentials with multi-scale brain parcellations and connectomes

**DOI:** 10.1101/2021.03.16.435599

**Authors:** David Pascucci, Sebastien Tourbier, Joan Rué-Queralt, Margherita Carboni, Patric Hagmann, Gijs Plomp

## Abstract

We describe the multimodal neuroimaging dataset VEPCON (OpenNeuro Dataset ds003505). It includes raw data and derivatives of high-density EEG, structural MRI, diffusion weighted images (DWI) and single-trial behavior (accuracy, reaction time). Visual evoked potentials (VEPs) were recorded while participants (n=20) discriminated briefly presented faces from scrambled faces, or coherently moving stimuli from incoherent ones. EEG and MRI were recorded separately from the same participants. The dataset contains pre-processed EEG of single trials in each condition, behavioral measures, structural MRIs, individual brain parcellations at 5 spatial resolutions (83 to 1015 regions), and the corresponding structural connectomes computed from fiber count, fiber density, average fractional anisotropy and mean diffusivity maps. For source imaging, VEPCON provides EEG inverse solutions based on individual anatomy, with Python and Matlab scripts to derive activity time-series in each brain region, for each parcellation level. The BIDS-compatible dataset can contribute to multimodal methods development, studying structure-function relations, and to unimodal optimization of source imaging and graph analyses, among many other possibilities.

## Background & Summary

Visual evoked potentials (VEPs) have a long record of shedding light on the spatial and temporal dynamics of large-scale neural processing in the brain^1,2^. EEG potentials registered at scalp electrodes result from synchronous activity in large populations of neurons that are distributed across cortical and subcortical areas^3,4^. Visual stimulation gives rise to a fast sequence of well-known EEG components that reflect initial processing at latencies before ∼100 ms^5,6^, subsequent object and recurrent processes^7–9^, and later components that reflect target detection, integration and decisions^10–12^. The VEP provides a millisecond by millisecond recording of whole-brain activity dynamics, and has a rich distribution of temporal frequencies that provides further insight into the functionality of brain processes^13,14^.

From VEPs recorded across the scalp, the underlying distributed patterns of brain activity can be estimated using an inverse solution based on anatomical constraints^15–18^. The anatomical constraints determine what electrical fields from a neural activity source would look like at the recording electrodes on the scalp, given the conductivities of various tissues and fluids lie in between. An inverse solution translates the recorded electrical potential field back to a pattern of distributed source activity in the brain. Source localizations from EEG are necessarily coarse, as compared to fMRI, and improving them further is an active field of research and helps understand the activity dynamics within areas, and their inter-relatedness^19–22^.

Activity in each area can influence activity throughout the brain in a few steps, due to the dense connectivity of cortico-cortical and cortico-subcortical fibers^23–26^. This structural connectivity can be inferred with diffusion weighted imaging (DWI)^27,28^. The resulting connectome constitutes a road map of sorts over which activity can propagate between areas, and in an important sense constrains how activity within an area can evolve through the influences it receives from others^29,30^.

The VEPCON dataset combines VEPs, T1-weighted (T1w) MRI, and DWI for 20 human participants, with as derivatives inverse solution matrices, brain parcellations and connectomes at 5 different spatial scales (Figure 1). These data were recorded to study the dynamics of functionally specialized processes that support face and motion perception. High-density EEG was recorded in two active paradigms where participants categorically discriminated face images from scrambled counterparts, or coherent from incoherent motion in random dot kinematograms. VEP sources for face stimuli are known to include inferior temporal and lateral occipital cortex and for motion stimuli they include dorsal area MT^8,9^. Part of these data were previously used for improving and validating EEG source imaging methods^31^ and time-varying functional connectivity methods^32,33^, for using connectomes to inform inverse solutions^34,35^, as well as for developing the multi-modal imaging platform Connectome Mapper 3^36^.

**Figure 1.**
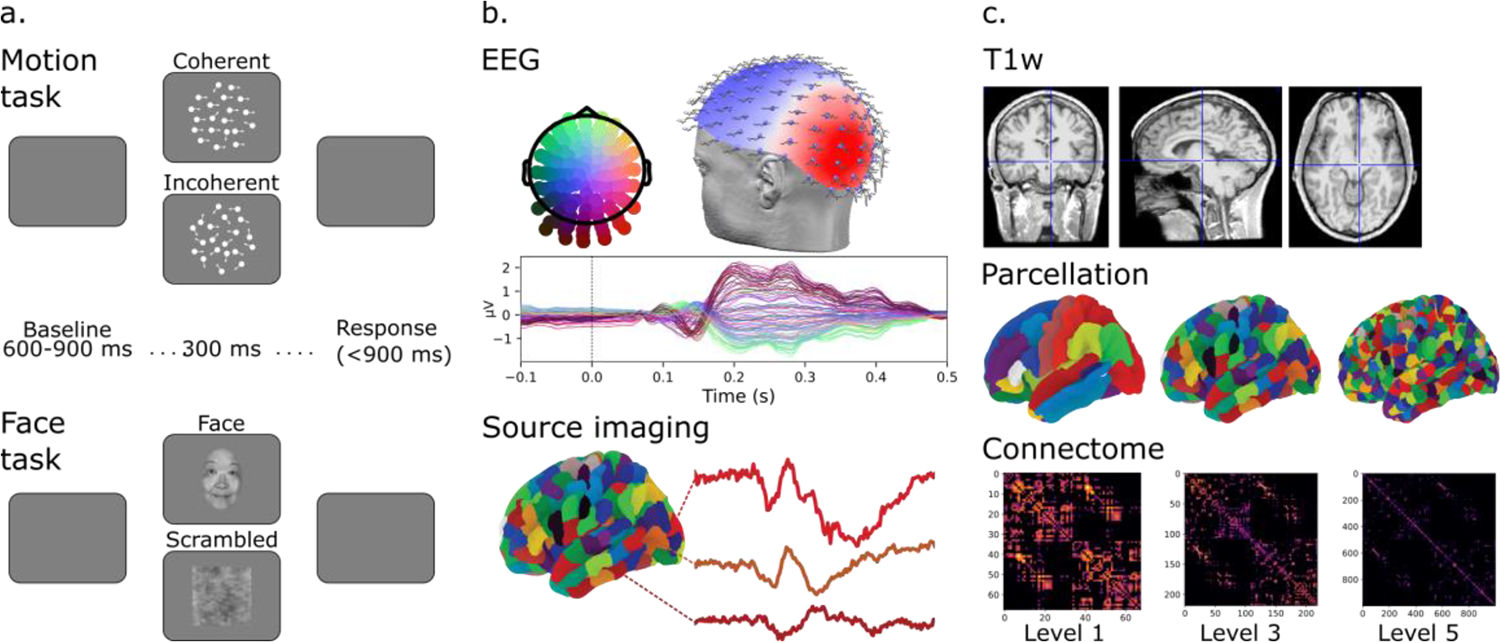
Schematic overview of the study design and data. a) Stimuli and temporal structure of trials in the Motion and Face task. b) Illustrates 128-channel EEG recording and a grand-average visual evoked potential at the scalp level, with corresponding source activity time-series from three areas at parcellation Level 3. c) Sample defaced T1w images and illustration of brain parcellations and corresponding connectomes based on fiber density at three spatial scales.

The dataset is publicly available as OpenNeuro Dataset ds003505, and structured following the MRI and EEG Brain Imaging Data Structure standards (BIDS)^37,38^. We expect the data to be useful for the development and benchmarking of multimodal analysis methods that combine functional and structural information, for exploring structure-function relations, and for controlling whether and how the level of parcellation affects results. We also expect unimodal reuse value for development and benchmarking of EEG source-imaging approaches and functional connectivity analyses. Beyond this, the availability within the same participant of single-trial behavior, EEG data, inverse solutions, anatomical and diffusion MRI, and connectomes based on four different indices allows for many types of analyses that can generate new hypotheses about dynamic brain function in human visual processing and behavior.

## Methods

### Participants

Twenty participants (3 males, mean age = 23 ± 3.5) took part that were recruited from the local student population (University of Fribourg, Switzerland). Participants had normal or corrected-to-normal vision. Before the experiment visual acuity was tested with the Freiburg Acuity test, and a value of 1 had to be reached with both eyes open ^39^. Nineteen participants were right handed, one was left handed. All participants provided written informed consent before the experiment. The experimental procedures complied with the Declaration of Helsinki and were approved by the regional ethics board (CER-VD, Protocol Nr. 2016-00060).

### Stimuli, display and procedures

EEG data were recorded while participants performed a face detection and a motion discrimination task (see Figure 1). The order of the two tasks was counterbalanced across participants.

Face stimuli were female and male faces (4° by 4° of visual angle) taken from online repositories and cropped with a Gaussian kernel to smooth the borders. Scrambled images were obtained by fully randomizing the phase spectra of the original images ^40^. In the face detection task, each trial lasted 1.2 s and started with a blank screen (500 ms). After the blank screen, one image (either a face or a scramble image of a face) was presented at the center of the screen for 200 ms and participants had the remaining 1000 ms to respond. The task was to report whether they saw a face or not (yes/no task) by pressing one of two buttons in a response box with their right hand. After the response and a random interval (from 600 to 900 ms), a new trial began. The experiment consisted of four blocks of 150 trials each, for a total of 600 trials, i.e., 300 with faces and 300 with scrambled faces. Faces and scrambled faces were randomly interleaved across trials.

In the motion discrimination task, motion stimuli were dot kinematograms presented on a circular frame at the center of the screen (dot field size = 8°; dot size = 12 pixels; lifetime = 10 frames; average number of dots inside the field = 75; dot speed in displacements per frame = 0.01^°;^ mean dot luminance =50%). For coherent motion, 80% of the dots were moving toward either the left or right, with the remaining 20% moving in random directions. For incoherent motion stimuli, all dots moved randomly. Each trial started with a blank interval of 500 ms followed by a centrally presented stimulus for 300 ms (Figure 1). Participants discriminated whether the presented motion was coherent or incoherent by pressing one of two buttons of a response box. After the response there was a random interval (from 600 to 900 ms) before the next trial began. There were four blocks of 150 trials each, for a total of 600 trials (300 with coherent motion). The two conditions were randomly intermixed within each block.

Stimuli were generated using Psychopy^41,42^ and presented on a VIEWPixx/3D display system (1920 × 1080 pixels, refresh rate of 100 Hz). Responses were collected using a ResponsePixx response box (VPixx technologies).

### EEG recording and preprocessing

EEG data were recorded at a sampling rate of 2048 Hz with a 128-channel Biosemi Active Two EEG system (Biosemi, Amsterdam, The Netherlands) in a dimly lit and electrically shielded room. Signal quality was ensured by monitoring and maintaining the offset between the active electrodes and the Common Mode Sense – Driven Right Leg (CMS-DRL) feedback loop under a standard value of ±20 mV. After each recording session, individual 3D electrode positions were digitized using an ultrasound motion capture system (Zebris Medical GmbH).

Offline, data were preprocessed using EEGLAB 14.1.1^43^. EEG data were downsampled to 250 Hz and detrended (antialiasing filter: cut-off = 112.5 Hz, bandwidth = 50 Hz, detrending at <1 Hz). Line noise (50 Hz) was removed via spectrum interpolation^44^. Data were then segmented into epochs and time locked from -1500 to 1000 ms from the stimulus onset in both tasks. Data from participant 05 (Face and Motion dataset) and 15 (Motion dataset) was not further processed due to excessive noise. Bad channels and epochs were identified and removed before preprocessing. Remaining physiological artifacts were isolated using an independent component analysis (ICA) decomposition (FastIca). Bad components were labelled by crossing the results of a machine-learning algorithm (MARA, Multiple Artifact Rejection Algorithm in EEGLAB) with the criterion of >90% of total variance explained and removed manually. Bad channels were then interpolated using the nearest-neighbour spline method and data were re-referenced to the average reference.

### EEG source imaging

EEG source imaging was performed using Cartool (v3.80)^45^ and custom-made scripts (see Matlab and Python examples in the code/directory). Source reconstruction was based on Cartool-segmented individual MRI T1w data, co-registered individual electrode positions, and the LORETA^46^ and LAURA^15^ algorithms (regularization = 6; spherical model with anatomical constraints, LSMAC). Leadfields were calculated for each of the around 5000 freely oriented dipoles while limiting the solution space to gray matter voxels^45^.

### MRI recording and processing

MR data from the same 20 subjects was acquired on a General Electrics Discovery MR750 3T MRI clinical scanner at the cantonal hospital in Fribourg, Switzerland, using a 32-channel head coil. The acquisition included anatomical T1-weighted images and DTI. T1-weighted images were acquired as rapid-gradient echo (MPRAGE) volumes using a COR FSPGR BRAVO pulse sequence (flip angle = 9^°;^ echo time = 2.8 ms; repetition time = 7300 ms; inversion time = 0.9 s, FOV = 220 mm, matrix size = 256×256, number of slices = 276, slice thickness = 1 mm, in-plane resolution = 0.9×0.9 mm^2^). DTI data were acquired with a spin echo single shot EPI pulse sequence and a diffusion sensitizing gradient set of 30 different directions and 5 diffusion-free B0 scans (echo time = 87 ms; repetition time = 8000 ms; interleaved slice order; b-weighting of 1000, FOV = 260 mm, matrix size = 128×128, number of slices = 60; slice thickness = 2.0 mm; slice spacing = 0.2 mm, in-plane resolution = 2.0×2.0 mm^2^).

Processing of all T1w and DTI data was performed using the Connectome Mapper v3.0.0-beta-RC1 pipelines^36^. All T1w scans were resampled to 1mm^3^ isotropic resolution from which gray and white matter were segmented using Freesurfer 6.0.1 ^47^, and parcellated into 83 cortical and subcortical areas^48^. The parcels were then further subdivided following the method proposed by^49^ into 129, 234, 463 and 1015 approximately equally sized parcels according to the Lausanne anatomical atlas. DTI data were corrected from motion and eddy current distortions using mcflirt and eddy_correct provided by FSL 5.0.9 and resampled to 1mm^3^ resolution using mrconvert from MRtrix 3.0.0-RC1. Diffusion directions per voxel were then reconstructed using the algorithm of constrained spherical deconvolution implemented in MRtrix 3.0.0-RC1^50^ with a maximal order of 4, enabling the estimation of multiple directions per voxel.

For sharing purposes, all raw T1w and DTI data were anonymized during BIDS conversion and all anatomical T1w data were de-identified by removing facial features using Quickshear^51^.

### Structural connectomes

For each participant, structural connectivity matrices were estimated from the reconstructed fiber orientation distribution (FOD) image using the SD_stream deterministic streamline tractography algorithm implemented in MRtrix 3.0.0-RC1^50^. Fiber streamline reconstruction started from seeds in the white-matter that were spatially random and the whole process completed when a number of 1M fiber streamlines were reconstructed. At each streamline step of 0.5mm, the local FOD was sampled, and from the current streamline tangent orientation, the orientation of the nearest FOD amplitude peak was estimated via a Newton optimization on the sphere. Fibers were stopped if a change in direction was greater than 45 degrees. Fibers with a length not in the 5mm to 200mm range were discarded. The streamline reconstruction process was complete when both ends of the fiber left the white matter mask. Then, for each scale, the parcellation was projected to the native DTI space after symmetric diffeomorphic co-registration between the T1w scan and the diffusion-free B0 using the Advanced Normalization Tools (ANTs) 2.2.0. Finally, connectivity matrices at 5 different spatial scales were built by considering all fiber streamlines connecting parcels according to the following connectivity measures: number of fibers, fiber density, average and median Fractional Anisotropy (FA), and Mean Diffusivity (MD).

## Data Records

The VEPCON dataset is available via the Open Neuro repository (doi: 10.18112/openneuro.ds003505.v1.0.1), and is fully BIDS compatible (v1.4.1). Below, we describe all data records following the directory structure.

The main directory contains for each participant the MRI T1w data and diffusion weighted images (DWI) files, with acquisition parameters as a JSON file. It also contains a JSON file detailing age and sex of each participant.

The derivatives/directory contains EEG derivatives obtained with EEGLAB (eeglab_) and Cartool (cartool_) in their respective folders. The output reports generated by mriqc (mriqc_), and anatomical and diffusion MRI derivatives are in the Connectome Mapper 3 (cmp_) folder, where the version of each software used is encoded in the folder name (e.g. cmp_v3.0.0-beta-RC1).

In the eeglab/directory, each participant’s folder contains the single trial epochs (.fdt, .set). The FACES files contain epochs for face stimuli and control stimuli, the MOTION files contain epochs for coherent and incoherent motion stimuli. A corresponding Matlab file contains a table with behavioral data for each single trial, listing stimulus specifiers, response made, response evaluation, reaction time and whether trial was discarded or not in preprocessing. Trials with behavioral errors or reaction times slower than 200 ms are marked as outliers. The same behavioral data is also included in the EEG epoch files. The files MOTION_preprocessing_summary.mat and FACE_preprocessing_summary.mat summarize the proportions of channels, epochs and ICA components removed during preprocessing for each participant.

The cartool/directory holds a text file (.xyz) with individual electrode positions in mm, co-registered to the participant’s head (x, y, z, Biosemi electrode name). The .spi file is a similar text file listing x, y, z coordinates and a text label for each sourcepoint for which an inverse solution was calculated. The inverse solution matrices for LAURA and LORETA ^15,46^ are in the respective .is files, they map observed patterns of EEG potentials to a distribution of 3D dipoles across source points. The rois/folder lists files that indicate which source points belong to what area (region of interest, ROI), according to each parcellation level, with a Cartool readable (.rois) and a Python readable version (.pickle.rois).

The cmp/directory contains anatomical, diffusion and structural connectivity data for each participant. All T1w-derived data are placed in the anat/folder, which includes the brain segmentations and parcellations in the native T1w space and in the native DTI space (_space-DWI_). All DTI-derived data are placed in the dwi/folder. It includes the preprocessed DTI, the FOD image and the final tractogram used to build the connectivity matrices. The connectivity/folder holds connectomes at each parcellation level, with a Matlab readable (.mat) and Python readable (.gpickle) version. Each connectome file contains the following metrics: number_of_fibers, fiber_density, mean_FA and median_FA, mean_MD and median_MD. The transformations from the native T1w space to the native DTI space applied to the parcellations are stored in the xfm/folder. The generated .pdf in the group folder visualizes each individual’s connectome using the number of fibers metric, for parcellation scale 1.

The code/directory lists example code with dependencies to derive time-series of activity per ROI from the preprocessed EEG data, and example code used for removing facial features from T1w images ^51^.

All outputs generated by MRIQC to support quality assessment of T1w data can be found in the mriqc/directory ^52^.

## Technical Validation

### Behavioral variables

Analysis of proportion correct and reaction times showed that participants behaved according to task instruction. In the face detection task the average accuracy was 97 ± 2% of correct responses, with mean reaction times of 501 ± 60 ms. Trials with behavioral errors or reaction times slower than 200 ms were marked as outliers (mean proportion of outliers 0.03 ± 0.02).

In the motion discrimination task, the average accuracy was 90 ± 14% and mean reaction times were 680 ± 90 ms. The mean proportion of outliers was 0.10 ± 0.14. One participant (number 16) inverted the response keys for several trials in the motion task, leading to an outlier accuracy value (36% of correct responses).

### VEPs

EEG data were visually inspected and noisy trials and channels were excluded from further analysis (see EEG recording and preprocessing). Remaining ocular, muscle and other artifacts were removed using ICA. The average proportion of channels removed across participants was 0.12 ± 0.07, range 0.02-0.28; mean proportion of epochs removed due to non-stereotyped artifacts, peristimulus eye blinks and eye movements: 0.04 ± 0.06, range 0.002-0.26); proportion of ICA components removed: 0.05 ± 0.03, range 0.01-0.15.

The dataset includes detailed summaries of preprocessing and data cleaning on the subject and single trial level (see Data Records).

Figure 2 shows the grand-average VEPs for the Face, Scrambled Face, Coherent Motion and Incoherent Motion conditions, showing robust evoked responses with components that conform the existing literature.

**Figure 2.**
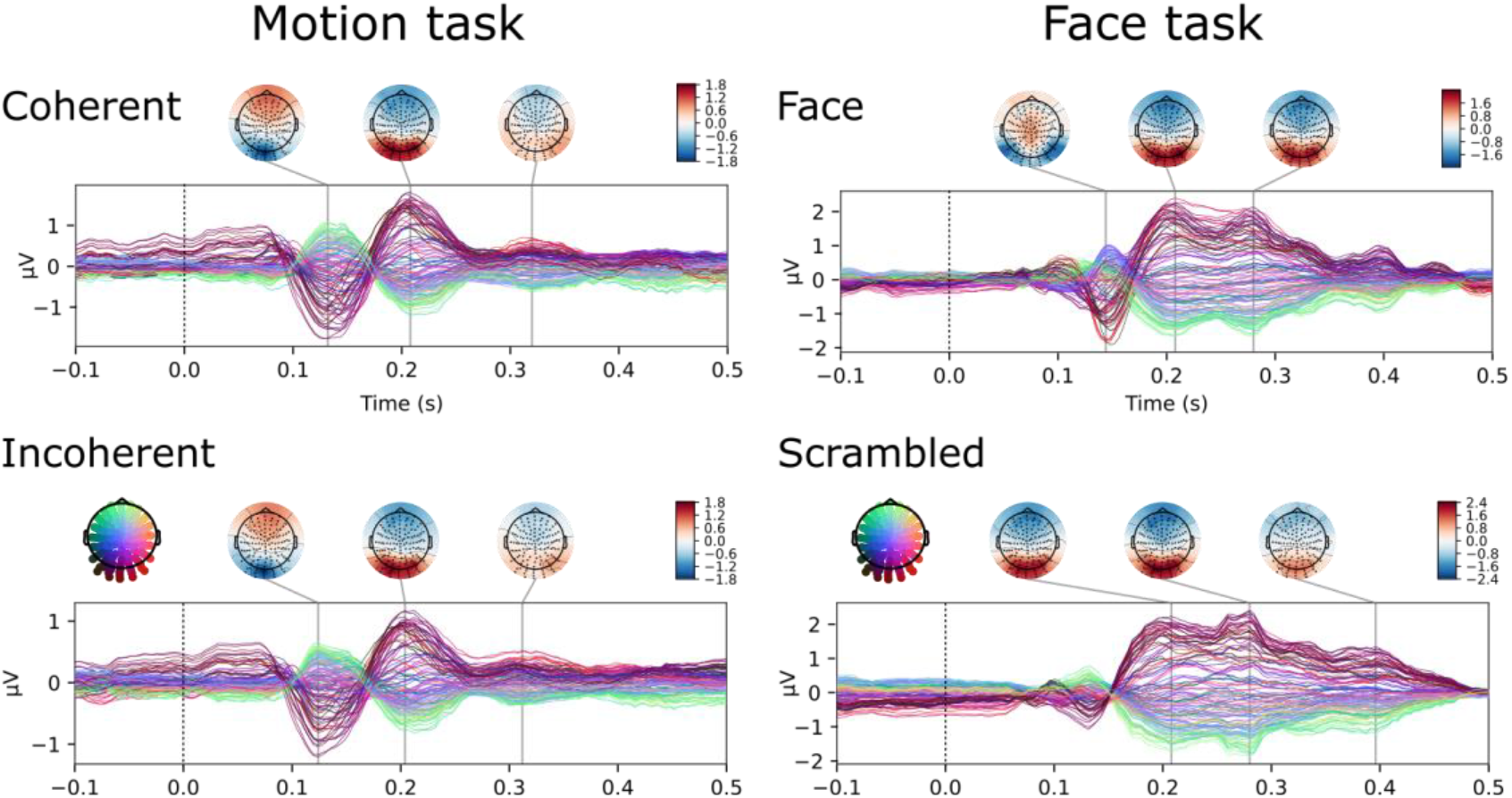
Grand-average VEPs for each of the four stimulus conditions, with corresponding scalp potential maps.

### MRI and connectomes

After recording MRI data were visually inspected and checked for neurological anomalies by the radiologist. Quality of T1w data was further inspected quantitatively by running MRIQC v0.16.1, an automated and robust quality control tool for T1w data which derives a set of 56 different image quality metrics to characterize image quality at different levels such as noise, motion or imaging artifacts ^52^. Results were summarized in reports at both individual and group levels, which are included in the mriqc/derivatives folder.

De-identification of T1w data was checked individually and supported by the creation of a PDF report that is available in the code/folder.

For each participant, the quality of the output after each processing step of the Connectome Mapper 3 pipelines was assessed using its graphical user interface, where we inspected individually the different Freesurfer outputs, brain parcellations, DTI data after preprocessing, co-registered T1w-derived parcellations with DTI data, the estimated FOD image, the reconstructed tractogram, and the reconstructed connectomes. Visual inspection did not reveal any artifacts.

## Usage Notes

For optimal sharing and re-usability, the dataset conforms to BIDS standards for MRI ^37^ and EEG ^38^.

Python and Matlab example code shows how to import the various files. This assures compatibility with commonly used EEG and MRI software, including MNE-Python ^53^, NeuroPycon ^54^, Fieldtrip ^55^ and SPM ^56^. The code also shows how to calculate one time-series of activity from all source points contained in a region using singular-value decomposition. Source activity for around 5000 freely oriented dipoles was extracted from all the source points inside each cortical area, as defined by the parcellation, and projected to a representative single direction using singular-value decomposition ^31^.

For creating new inverse solutions, new forward models can be created from the T1w images, using the individual electrode coordinate files. When new leadfields are generated using the defaced MRIs, small differences can occur, but are unlikely to pose problems for the intended reuse purposes ^57^.

## Code Availability

The dataset contains a code/folder with scripts, and .ini files for generating each derivative. Connectome Mapper 3 is freely available at https://connectome-mapper-3.readthedocs.io, the Cartool software via https://sites.google.com/site/cartoolcommunity.

## Acknowledgements

This research was supported by Swiss National Science Foundation grants PP00P1_183714, PP00P1_190065 and CRSII5-170873.

## Author contributions

Conceptualization GP, DP

Investigation DP

Formal analysis DP, ST, JRQ, MC

Data curation and Software ST, DP, JRQ

Supervision and Funding acquisition GP, PH

Writing original draft GP, DP, ST

Writing review & editing All authors

### Competing interests

All authors declare that they have no conflicts of interest in publishing this work and the associated data.

## Notes

https://openneuro.org/datasets/ds003505/versions/1.0.1

### Competing Interest Statement

The authors have declared no competing interest.

